# Action enhances predicted touch

**DOI:** 10.1101/2020.03.26.007559

**Authors:** Emily R. Thomas, Daniel Yon, Floris P. de Lange, Clare Press

## Abstract

It is widely believed that predicted tactile action outcomes are perceptually attenuated. The present experiments determined whether predictive mechanisms always generate attenuation, or instead can enhance perception – as typically observed in sensory cognition domains outside of action. We manipulated probabilistic expectations in a paradigm often used to demonstrate tactile attenuation. Participants produced actions and subsequently rated the intensity of forces on a passive finger. Experiment 1 confirmed previous findings that action outcomes are perceived less intensely than passive stimulation, but demonstrated more intense perception when active finger stimulation was removed. Experiments 2 and 3 manipulated prediction explicitly and found that expected touch during action is perceived *more* intensely than unexpected touch. Computational modelling suggested that expectations increase the gain afforded to expected tactile signals. These findings challenge a central tenet of prominent motor control theories and demonstrate that sensorimotor predictions do not exhibit a qualitatively distinct influence on tactile perception.

**Statement of Relevance:** Perception of expected action outcomes is thought to be attenuated. Such a mechanism may be adaptive because surprising inputs are more useful - e.g., signalling the need to take new courses of action - and is thought to explain why we cannot tickle ourselves and unusual aspects of action and awareness in clinical populations. However, theories outside of action purport that predicted events are perceptually facilitated, allowing us to generate largely accurate representations of our noisy sensory world. We do not know whether action predictions really alter perception differently from other predictions because different manipulations have been performed. Here we perform similar manipulations and demonstrate that action predictions can enhance, rather than attenuate, touch. We thereby demonstrate that action predictions may not have a qualitatively distinct influence on perception, such that we must re-examine theories concerning how predictions influence perception across domains and clinical theories based upon their assumptions.

---

When we produce actions we predict their sensory consequences. Prominent motor theories (Blakemore et al., 1998; Dogge et al., 2019; Fiehler et al., 2019; Kilteni & Ehrsson, 2017) propose that we attenuate – or downweight – perception of expected action outcomes. Such downweighting mechanisms are thought to finesse the limited capacity of our sensory systems, prioritising perception of more informative unexpected sensory events that signal the need to perform new actions or update of our models of the world (Press et al., 2020; Wolpert & Flanagan, 2001). For example, if we lift a cup of coffee that is lighter than expected, attenuated processing of expected signals (e.g., touch on our fingertips) will prioritise perception of unexpected events (e.g., accelerating motion of the cup) allowing swift updating of our beliefs about the environment (e.g., the weight of the cup) and support corrective action to avoid spillage. These downweighting mechanisms are invoked to explain that self-produced tactile sensations generate lower activity in bilateral secondary somatosensory cortex (Blakemore et al., 1998; Kilteni & Ehrsson, 2020; Shergill et al., 2013), and are perceived to be less intense (Bays et al., 2005, 2006; Kilteni et al., 2019; Shergill et al., 2003; Wolpe et al., 2016, 2018), than externally-produced forces. This theory also provides an explanation for why it is difficult to tickle oneself (Blakemore et al., 1998).

However, outside of action it is thought that prediction mechanisms generate a qualitatively opposite influence on perception. In these theories – typically couched in Bayesian frameworks – it is proposed that we combine our expectations (prior) with the input (likelihood) to determine what we perceive (posterior; de Lange et al., 2018). Such a process would *upweight*, rather than downweight, perception of expected events, increasing the detectability and apparent intensity of events (Brown et al., 2013; Carrasco et al., 2004) and thereby enabling rapid generation of largely veridical experiences in the face of sensory noise (de Lange et al., 2018; Kersten et al., 2004). For example, some theories propose that when we predict a sensory event we increase the gain on sensory units tuned to that event and via competitive local interactions relatively inhibit sensory populations tuned to unpredicted events (de Lange et al., 2018; Kok et al., 2012; Press et al., 2020; Press & Yon, 2019; Summerfield & de Lange, 2014; Yon et al., 2018). It is perhaps unclear why the adaptive arguments presented for downweighting (informativeness) and upweighting (veridicality) predicted perceptual experiences should apply differentially in the domain of action, and some evidence from the visual domain suggests that they may not (Yon et al., 2018, 2020; Yon & Press, 2017).

Interestingly, a stark difference between studies purporting to demonstrate upweighting and downweighting is that the former study visual perception whereas the latter study tactile perception. It is therefore widely believed that action predictions shape tactile perception in a qualitatively distinct way, including proposals that differences relate to tactile events being body-related (Dogge et al., 2019) and tightly coupled with the motor system (Kusnir et al., 2019), in a way that many predicted visual or auditory events are not. Similarly, differences may also relate to assumptions that tactile attenuation during action is dependent upon somatosensory-cerebellar connectivity (Blakemore et al., 1998; Kilteni & Ehrsson, 2020), in contrast with hippocampal mediation of prediction in visual processing (Kok et al., 2020; Kok & Turk-Browne, 2018).

However, studies examining touch perception during action have not manipulated predictability in the same way as those examining the wider influences of prediction on perception and therefore direct comparisons are difficult. The defining feature of prediction mechanisms is that they operate according to stimulus probabilities (de Lange et al., 2018). As such, prediction mechanisms outside of action contexts are typically measured by presenting events with high and low conditional probabilities, and comparing perception of the ‘expected’ with the ‘unexpected’. In contrast, typical studies demonstrating tactile attenuation during action compare the perception of events in the presence or absence of action, or when events are coincident versus delayed with respect to action (Bays et al., 2005, 2006; Blakemore et al., 1998; Kilteni et al., 2019; Shergill et al., 2013; Wolpe et al., 2018). Thus, manipulating conditional probabilities is essential for establishing comparability between domains, and thereby whether action predictions really influence touch perception via qualitatively distinct mechanisms from other types of prediction. To address this question the present studies adopted a force judgement paradigm used widely in action domains to examine tactile attenuation, and additionally employed probabilistic predictive manipulations.

## General Method

### Participants

Thirty distinct participants were tested in Experiment 1 (16 female, mean age = 25.53 years [SD = 5.25]), Experiment 2 (20 female, mean age = 22.80 years [SD = 3.18]) and Experiment 3 (22 female, mean age = 24.3 years [SD = 4.34]). Eight participants in Experiment 1, six participants in Experiment 2, and nine participants in Experiment 3, were replacements for those where acceptable psychometric functions could not be modelled to their responses (flat functions), where they were unable to follow instructions concerning movement performance (>20% recorded movement errors), or where there was technical malfunction. These criteria were established a priori to participant testing and replacements resulted in a total sample of 30 participants in each experiment. One participant’s PSE score from Experiment 2 was winsorized to meet the normality assumptions of parametric tests (from z = 3.34 to z = 3, Tukey, 1962). Participants were recruited from Birkbeck, University of London and paid a small honorarium for their participation. All participants reported no current neurological or psychiatric illness and provided written informed consent prior to participation. The experiments were performed with local ethical committee approval (Birkbeck, University of London) and in accordance with the ethical standards laid down in the 1964 Declaration of Helsinki. The sample size was determined a priori on the basis of pilot testing to estimate effect size – to have at least 80% power of detecting medium effect sizes (*d* = 0.5) – and parametric assumptions were met.

#### Experiment 1

The first aim of Experiment 1 was to determine whether we could replicate typical action attenuation effects within our set-up. Participants therefore moved an active right index finger towards a passive left index finger, similarly to in typical force judgement tasks (Bays et al., 2005, 2006; Kilteni et al., 2019) – where passive finger touch is reported as less forceful during action than when the active finger remains still. Our ‘Contact’ condition closely resembled these previous studies (see Fig. 1A). We were also interested in whether similar effects would be found in a ‘No Contact’ condition, where an identical downward motion of the active finger triggered the same stimulation but without making contact with any device. This condition was included to investigate the nature of effects before manipulating conditional probabilities, because if sensorimotor prediction is underpinned by predictive mechanisms defined according to event probabilities then active-finger stimulation should not be required per se. Additionally, tactile attenuation could in principle be generated by generalised sensory gating mechanisms when there is active contact. Generalised gating is thought to reduce perception of any events delivered to moving effectors (Williams et al., 1998; Williams & Chapman, 2000), and therefore regardless of prediction, because it is thought to occur at the earliest relay in the spinal cord (Seki & Fetz, 2012). This is why studies seeking to investigate predictive sensory attenuation have examined perception on passive effectors (Bays et al., 2006). However, perception of concurrent gated stimulation on an active finger could still, in principle, bias responses about stimulation on the passive finger due to response biases (Firestone & Scholl, 2016).

**Figure 1.**
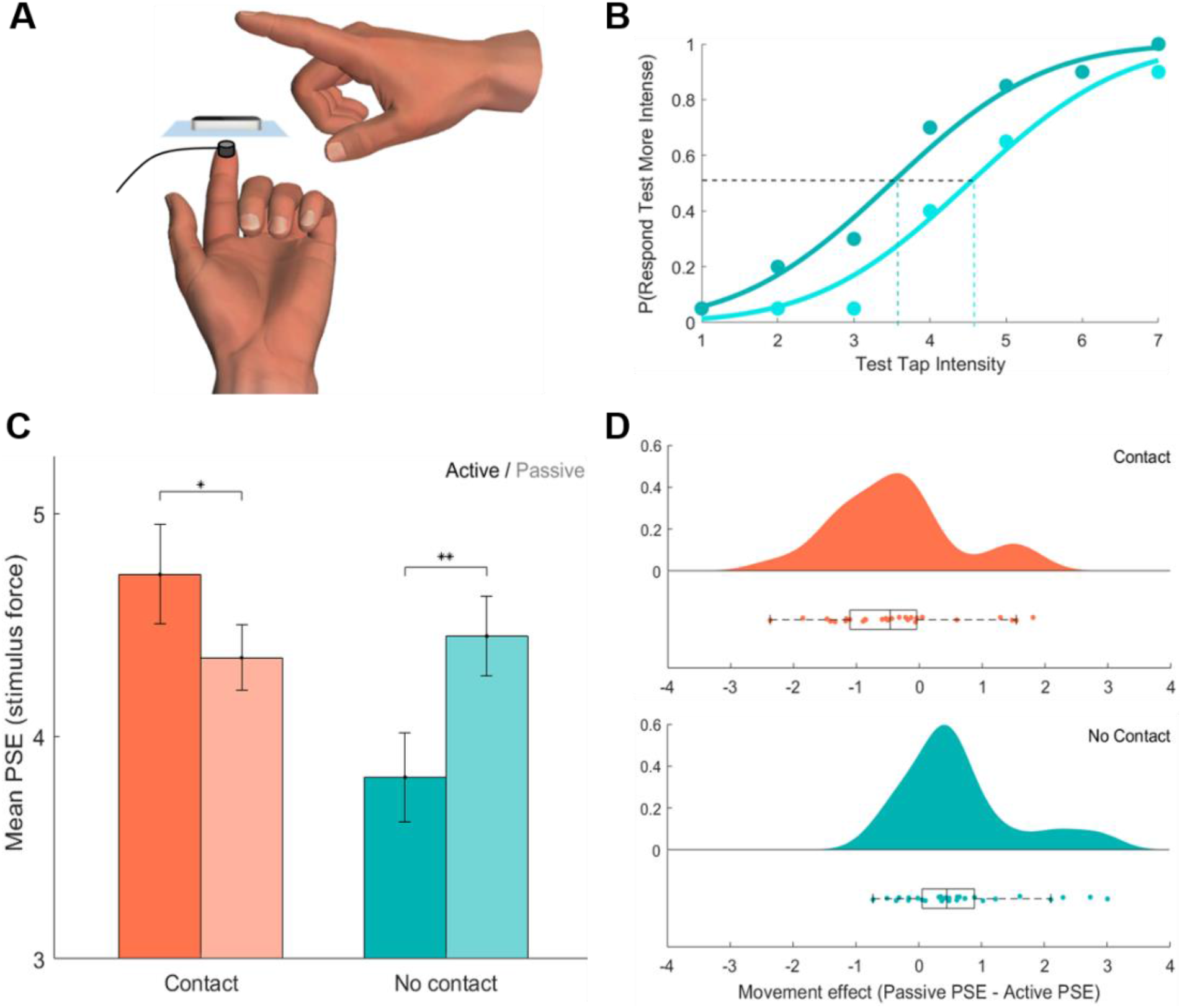
Experiment 1. (A) On each trial, participants made downward movements with their right index finger either over a motion tracker (No Contact condition), or towards a button with which they made contact (Contact condition). Each movement elicited a tactile punctate event to the left index finger positioned directly below. (B) PSEs were calculated for each participant (data represents an example participant in the No Contact condition for Active [dark blue] and Passive [light blue] trials). (C) Mean PSEs (± SEM) were higher in Active than Passive trials in the Contact conditions, but lower in Active than Passive trials in the No Contact condition. Larger PSEs indicate less intense target percepts (* *p* < .05, ** *p* = .001). (D) PSE effect of movement (Passive – Active) for the Contact (top) and No Contact (bottom) condition, plotted with raincloud plots displaying probability density estimates (upper) and box and scatter plots (lower). Boxes denote lower, middle and upper quartiles, whiskers denote 1.5 interquartile range, and dots denote difference scores for each participant (N=30). Positive effects of movement indicate more intensely perceived active events relative to passive events, but negative values indicate the reverse – less intensely perceived active events.

### Procedure

The experiment was conducted in MATLAB using the Cogent toolbox. Participants held their left hand palm upwards (Fig. 1A) with their index finger positioned against a solenoid (diameter of metal rod = 4 mm; diameter of solenoid = 15 mm; TACT-CONTR-TR2, Heijo Research Electronics) sitting on the apex of the fingertip. Their right hand rested on a shelf, positioned such that the index finger distal phalange was directly above the left hand distal phalange, but rotated 90 degrees anticlockwise relative to their left hand (Fig 1A). An infrared motion tracker (Leap Motion Controller using the Matleap MATLAB interface) was placed on the shelf supporting the solenoids at the midpoint between them.

At the start of each trial, participants were cued onscreen to move their right index finger (‘move’; Active trials – 50%) or remain stationary (‘do not move’; Passive trials – 50%). On Active trials, they rotated their index finger downwards at the metacarpophalangeal joint. When motion was detected – by either a button press in Contact blocks or infrared tracking in No Contact blocks – the target stimulus was delivered to the left index finger for 30 ms, resulting in apparent synchrony of stimulation with movement. After 1000 ms, a reference stimulus was presented for 30 ms. The target stimulus presented one of seven logarithmically-spaced forces, and the reference stimulus always presented the fourth (middle) force. After a 300 – 500 ms delay, participants were asked which tap was more forceful, responding with a left foot pedal for the first stimulus and a right foot pedal for the second stimulus. The next trial started after 1000 ms. In Passive trials, the target stimulus was delivered 500 ms after the cue to remain still. Participants’ hands were visually occluded during the experiment and white noise was played through headphones (53 Db; piloting confirmed that this level resulted in inaudible solenoid movement) throughout testing.

In Contact blocks, participants’ right hand was positioned 5 cm above their left hand, and in No Contact blocks it was moved to 12 cm above to allow movements to be made without touching the shelf. Termination points of movements were approximately the same in Contact and No Contact conditions. It is worth noting that this palm separation generated a difference between our Contact and No Contact conditions additionally to contact. This allowed us to replicate the typical setup in the Contact condition while allowing movement to be registered with the infrared tracking in the No Contact condition. Importantly, conclusions relate primarily to the simple effects within the Contact and No Contact conditions, so this additional difference should not alter the conclusions.

There were 560 trials in total; 140 for each of the Active and Passive conditions, in both the Contact and No Contact blocks. The order of blocks was counterbalanced across participants and trial type order was randomized. Participants completed eight practice trials before the main test blocks.

### Modelling Psychometric Functions

Participant responses were modelled by cumulative Gaussians to estimate psychometric functions, using the Palamedes Toolbox (Prins & Kingdom, 2018) in MATLAB. This procedure was performed separately for active and passive trials during the test phase. The mean of the modelled Gaussian was taken as the Point of Subjective Equivalence (PSE), describing the point at which participants judge the target and reference events to have equal force. Lower values are indicative of more intense target percepts.

### Results

PSE values were analysed in a 2×2 within-participants ANOVA, revealing no main effect of Contact (*F*(1, 29) = 3.11, *p* = .089 *ηp^2^* = .10) or Movement (*F*(1, 29) = 1.24, *p* = .274, *ηp^2^* = .04). However, there was a significant interaction between Contact and Movement (*F*(1, 29) = 15.39, *p* < .001, *ηp^2^* = .35), driven by lower force judgements (higher PSEs) in Active (*M* = 4.73, *SD* = 1.22) compared to Passive trials (*M* = 4.35, *SD* = .80) in the Contact condition (*t*(29) = 2.07, *p* = .047, *d* = .38), but higher force judgements (lower PSEs) in Active (*M* = 3.82, *SD* = 1.11) compared to Passive trials (*M* = 4.45, *SD* = .97) in the No Contact condition (*t*(29) = −3.80, *p* = .001, *d* = .69, see Fig. 1C).

#### Experiment 2

Experiment 1 revealed that when a moving effector receives cutaneous stimulation simultaneously with passive effector stimulation, tactile events are perceived less intensely during movement. Conversely, when the moving effector does not receive stimulation, tactile events are perceived more intensely during movement. These results replicate previous findings (Bays et al., 2005, 2006; Kilteni et al., 2019) that tactile events on a passive finger are perceived less intensely during active movement, but only when the specifics of the paradigm are replicated such that the active finger is also stimulated. One possible explanation of the difference between conditions is that generalised gating contributes to effects in the Contact condition. Due to this possibility, Experiment 2 examined the nature of the enhancement effect observed in the No Contact condition. Mechanisms generating this effect may operate according to predictive upweighting mechanisms, given that the tactile event can be anticipated on the basis of action, however its nature is difficult to determine without manipulating conditional probabilities between actions and outcomes (de Lange et al., 2018).

Therefore, in Experiment 2 we compared perception of tactile events when they were expected or unexpected based on learned action-outcome probabilities established in a preceding training session. Comparing expected and unexpected conditions – rather than the typical active vs passive comparison – allows us to isolate influences of prediction on tactile perception, while controlling for non-predictive influences of action on perception (Press & Cook, 2015). Downweighting accounts predict that participants will rate expected events as less intense (forceful), whereas upweighting theories predict that expected tactile stimulation will be rated more intensely, than unexpected stimulation.

### Procedure changes relative to Experiment 1

Participants now performed one of two movements that predicted one of two tactile effects (Fig. 2A). Participants were positioned with their left index and middle finger making contact with independent solenoids (Fig. 2A). At the beginning of each trial an arbitrary cue (either a square or circle) instructed participants to move their right index finger either upwards or downwards from the metacarpophalangeal joint, tracked by an infrared motion sensor. This action triggered delivery of the target stimulus to either solenoid. Presenting two action types and two stimulation types allowed us to compare perception of expected and unexpected events, while controlling for repetition effects. It should be noted in explicating the logic of the procedure here that any action predictions should determine *where* stimulation will be received, rather than its intensity. However, in Bayesian models it is assumed that enhanced detection and intensity of expected events relates to the precision of the estimate (Brown et al., 2013, e.g., a force precisely estimated to have occurred on a certain region of tactile space should be feel more intense because of the precise estimate of spatial information, rather than an estimate of the force per se). Since these models assume that predictions enhance the precision of resultant estimates, they would also predict enhancements in perceived force (and indeed other sensory attributes, like brightness or loudness).

**Figure 2.**
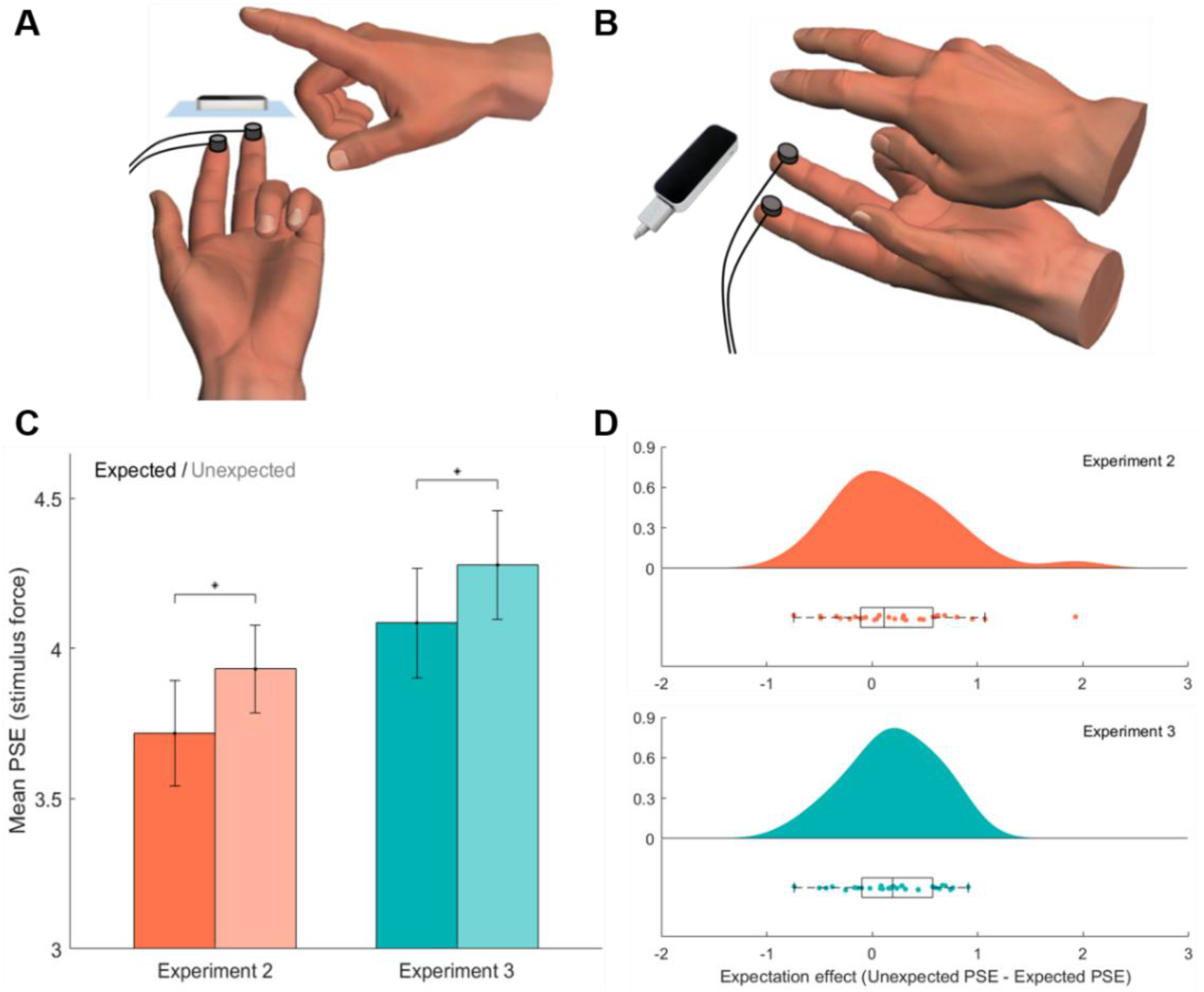
(A) Experiment 2. On each trial, participants made a downwards or upwards movement with their right index finger over a motion tracker, which elicited tactile punctate events to the left index or middle finger. Movements were perfectly predictive of tactile events during the training session, and 66.6% predictive during the test session. (B) The design of Experiment 3 was similar to Experiment 2, but participants instead made only downwards movements, now with either their right index or middle finger. (C) Mean PSEs (± SEM) were lower for expected than unexpected trials in both Experiment 2 and Experiment 3. Larger PSEs indicate a less intensely perceived target stimulus (* p < .05). (D) PSE expectation effect (Unexpected – Expected) plotted with raincloud plots displaying probability density estimates (upper) and box and scatter plots (lower), for Experiment 2 (top) and Experiment 3 (bottom). Boxes denote lower, middle and upper quartiles, whiskers denote 1.5 interquartile range, and dots denote difference scores for each participant (N=30). Positive expectation effect values indicate more intensely perceived expected events relative to unexpected events.

During training, participants’ action (e.g. downwards movement) was 100% predictive of the location of tactile event (e.g. left index finger). In a test session 24 hours later, the action-outcome relationship was degraded to measure perception of expect and unexpected events – the expected finger was stimulated on 66.6% of trials, and the unexpected was stimulated on the remaining 33.3% of trials. The training and test sessions were carried out at the same time on consecutive days. There were 420 trials in each session. Trial order was randomised and the action-stimulus mapping was counterbalanced across participants. The cue-action mapping was reversed half way through each session to account for effects resultant from possible learning of cue-outcome associations instead of action-outcome associations. Participants completed 12 practice trials before the main session trials.

### Results

Psychometric functions were modelled to participants responses similarly to in Experiment 1, but now separately for expected and unexpected events. PSE values were lower on expected trials (*M* = 3.72, *SD* = .96) than unexpected trials (*M* = 3.93, *SD* = .80; *t*(29) = −2.13, *p* = .041, *d* = .39, see Fig. 2C), demonstrating more forceful perception of expected than unexpected action outcomes.

#### Experiment 3

Experiment 3 was designed to provide a conceptual replication of Experiment 2, using a set-up more closely aligned with typical action paradigms – whereby one always makes a movement towards another effector. We additionally controlled for possibilities that the expectation effect in Experiment 2 may have resulted from cue-outcome learning by removing the cue stimulus and requiring free selection of action. The explicit reference stimulus was also removed and comparisons were made against an implicit reference, eliminating the possibility that effects are determined by forming predictions about the reference stimulus.

### Procedure changes relative to Experiment 2

Independent solenoids were now attached to both fingers via adhesive tape (diameter of metal rod = 4 mm; diameter of solenoid = 18 mm; TactAmp 4.2 Dancer Design). The foot pedals were positioned at either 45 (for stronger) or 90 (for weaker) degree angles relative to their right foot to record responses. At the start of each trial, participants selected to make a downwards movement with their right index or middle finger. Participants’ hands, and therefore index and middle fingers, were spatially aligned with each other (Fig. 2B). Actions were freely selected and the frequency of index and middle finger movements was monitored to ensure approximately equal numbers of both action types. Participants’ actions (e.g. downwards movement) were still perfectly predictive of the location of tactile events (e.g. left index finger) during training, and this contingency was again degraded to 66.6% in the following test session. Participants were asked whether they perceived the test force to be more or less forceful than the average force intensity. An example of the average force was presented to each finger once every 21 trials (NB: the average force was identical to the reference force intensity in Experiments 1 & 2). The experiment consisted of two training blocks followed by a test block, all occurring in the same session of testing. In the first training block participants responded yes/no to the question ‘Tap on index or middle finger?’, and in the second training block they were asked about the force, similarly to in Experiment 2 and in subsequent test blocks. Half of the participants experienced a mapping whereby moving the right hand index finger resulted in left hand index stimulation and middle finger movement resulted in middle stimulation. The other half experienced a mapping whereby index finger movement resulted in middle stimulation, and middle finger movement in index stimulation. There were 210 trials in each session.

### Results

Like in Experiment 2, PSE values were lower in expected (*M* = 4.08, *SD* = 1.00) than unexpected (*M* = 4.28, *SD* = .99) trials (*t*(29) = −2.56, *p* = .016, *d* = .47, see Fig. 2C), again demonstrating more forceful perception of expected than unexpected action outcomes.

## Computational modelling

The present findings are consistent with predictive upweighting theories of perception, which propose that it is adaptive for observers to combine sampled sensory evidence with prior knowledge – biasing perception towards what we expect (Kersten et al., 2004). This may be achieved mechanistically by altering the weights on sensory channels, increasing the gain of expected relative to unexpected signals (de Lange et al., 2018; Summerfield & de Lange, 2014). However, an alternative explanation is that expectation effects reflect biasing in response-generation circuits – such that action biases people to *respond* that events are more intense when they are expected, rather than altering perception itself (Firestone & Scholl, 2016). These different kinds of bias can be dissociated in computational models that conceptualise perceptual decisions as a process of evidence accumulation.

### Drift Diffusion Modelling procedure

Perceptual biases are modelled as growing across time – every time response units sample from perceptual units they will be sampling from a biased representation, therefore increasing the magnitude of biasing effects across a larger number of samples (Yon et al., 2020). In contrast, response biases are modelled as operating regardless of current incoming evidence and to be present from the outset of a trial (Leite & Ratcliff, 2011). According to this logic, we can model the decision process with drift diffusion modelling (DDM; Ratcliff & McKoon, 2008) to identify the nature of the biasing process. We can thus establish whether action expectations shift the starting point of evidence accumulation towards a response boundary (‘start biasing’, *z* parameter; Fig. 3A), or instead bias the rate of evidence accumulation (*db* parameter, ‘drift biasing’, Fig. 3B).

**Figure 3.**
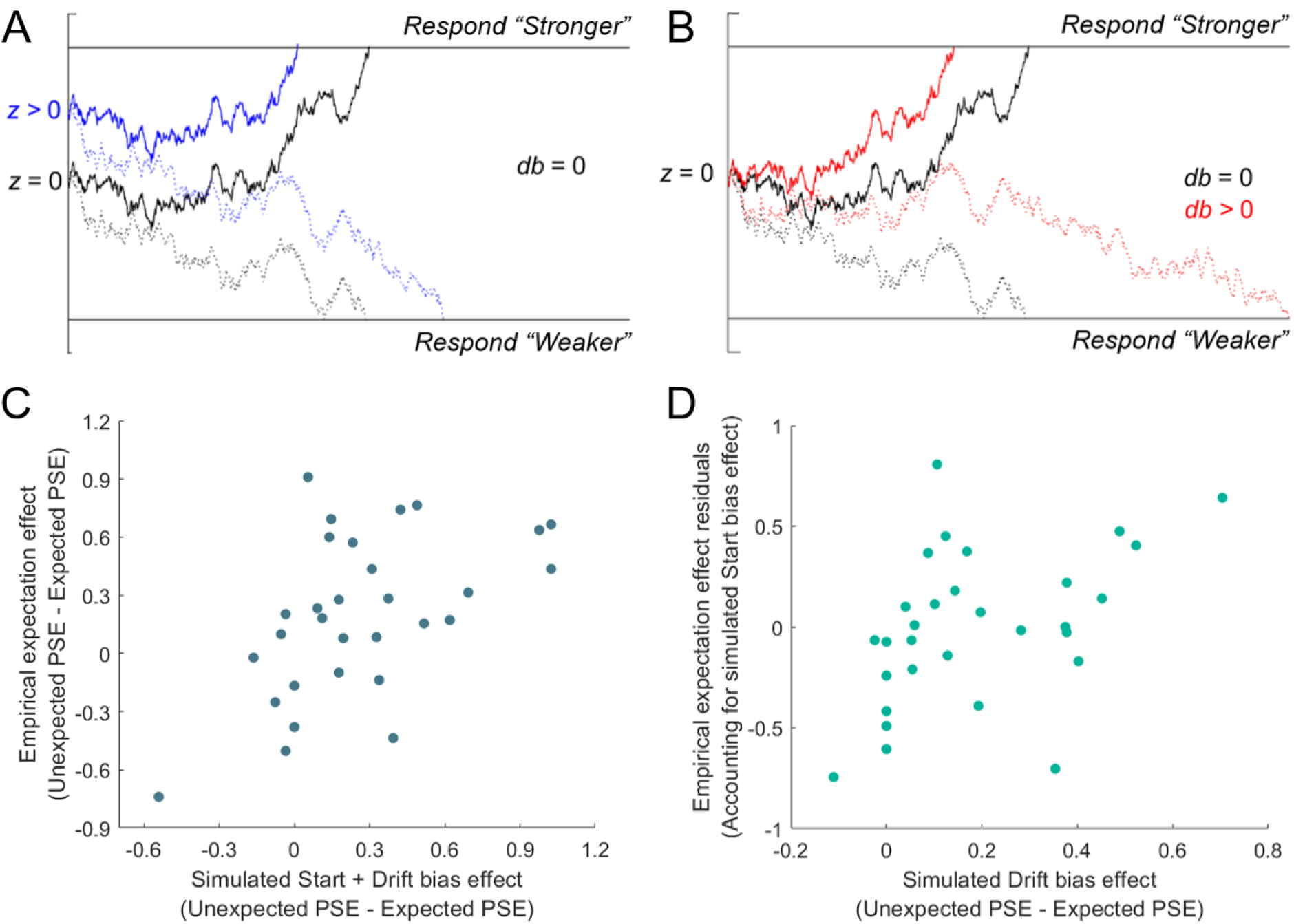
Illustration of how the DDM could explain expectation biases, and results of computational modelling. (A) For an unbiased decision process (black lines), sensory evidence integrates towards the upper response boundary when stimuli are stronger than average (solid lines) and towards the lower response boundary when weaker than average (dotted lines). Baseline shifts in decision circuits could shift the start point of the accumulation process nearer to the upper boundary for expected events (influencing the parameter *z*; blue lines - Start bias model). (B) Alternatively, selectively altering the weights on sensory channels could bias evidence accumulation in line with expectations (influencing parameter *db*; red lines – Drift bias model). (C) Simulated Start + Drift bias (winning DIC model) expectation effect plotted against the empirical expectation effect, showing a significant positive correlation. (D) Simulated Drift bias expectation effect plotted against the empirical expectation effect accounting for simulated Start bias effects (plotted as the residuals from a model where the simulated Start bias effect predicts the empirical effect), again showing a significant positive correlation. Importantly, our regression analysis revealed that drift biases accounted for significant additional variance once accounting for start biases. All expectation effects were calculated by subtracting Expected PSEs from the Unexpected PSEs.

We fit drift diffusion models to participant choice and reaction time data from Experiment 3 using the hDDM package implemented in Python (Wiecki et al., 2013; NB: reaction times were not collected in Experiment 2). In the hDDM, model parameters for each participant are treated as random effects drawn from group-level distributions, and Bayesian Markov Chain Monte Carlo (MCMC) sampling is used to estimate group and participant level parameters simultaneously. We specified four different models: 1) a *null* model where no parameters were permitted to vary between expected and unexpected trials; 2) a *start bias* model where the start point of evidence accumulation (*z*) could vary between expectation conditions; 3) a *drift bias* model where a constant added to evidence accumulation (*db*) could vary according to expectation; 4) a *start + drift bias* model where both parameters could vary according to expectation.

All models were estimated with MCMC sampling, and parameters were estimated with 30,000 samples (‘burn-in’=7,500). Model convergence was assessed by inspecting chain posteriors and simulating reaction time distributions for each participant. Models were compared using deviance information criteria (DIC) as an approximation of Bayesian model evidence, a common method used to determine model fit. Lower DIC values relative to a baseline, or null, model are indicative of a better model fit.

A posterior predictive check was conducted using the hDDM package to establish how well each model was able to reproduce the patterns in our data. The posterior model parameters for the *start bias*, *drift bias*, and *start + drift bias* models were used to simulate a distribution of 500 reaction times and choices for each trial for each participant. From this simulated data we calculated the probability that a ‘stronger than average’ response was given at each intensity level, separately for expected and unexpected trials. This allowed us to model simulated psychometric functions for expected and unexpected trials, exactly as we had done for empirical decisions. Performing this procedure for each model yielded separate simulated expectation effects (Unexpected PSE – Expected PSE) for each participant under the *start bias*, *drift bias* and *start + drift bias* models.

## Results

Fitting the DDM to the behavioural data found that the model allowing both start and drift biases to vary according to expectation provided the best fit (DIC relative to null = −234.8) relative to both the start bias (DIC relative to null = −191.06) and drift bias (DIC relative to null = −8.62) models. This finding may suggest that observed biases are a product of both start and drift rate biasing. However, although the DIC measure does include a penalty for model complexity, it is thought to be biased towards models with higher complexity (Wiecki et al., 2013) and it indeed favoured the most complex model here.

We conducted a posterior predictive check to evaluate how well simulated data from each of the models could reproduce key patterns in our data. Correlations were calculated to quantify how well simulated expectation effects reproduced empirical expectation effects, which revealed significant relationships for all three models (Start bias model: *r*_30_ = .39, *p* = .034; Drift bias model: *r*_30_ = .43, *p* = .017; Start + Drift bias model: *r*_30_ = .53, *p* = .003, see Fig. 3C).

Given that we were interested in whether any of the PSE expectation effect is generated by sensory biasing – rather than possible additional contributions of response biasing – we examined whether drift biasing accounted for any further variance in expectation effects than start biasing alone, by conducting a stepwise linear regression to predict the empirical expectation effect (Unexpected PSE – Expected PSE). In the first step, we included the simulated expectation effect from the start bias model to predict the empirical expectation effect. The simulated start bias data was able to predict the empirical expectation effect (*R*^2^ = .15, *F*(1,28) = 4.96, *p* = .034). In the second step, we included the simulated expectation effect from the drift bias model as an additional predictor of the empirical expectation effect, importantly providing a significant improvement to the model fit (*Fchange*(1,27) = 6.72, *p* = .015; *R*^2^ = .32, *F*(2,27) = 6.34, *p* = .006). This analysis reveals that a model implementing a drift biasing mechanism better predicts empirical effects of expectation on perceptual decisions, by explaining unique variance in participant decisions that cannot be explained by response biasing.

## Discussion

Extant models disagree about how predictions should shape perception of action outcomes. We examined whether sensorimotor prediction always attenuates perception of tactile events, as is widely assumed in the action literature, or whether predicted events may be perceptually upweighted – in line with theories in the wider sensory cognition literature. We adapted the force judgement paradigm typically used in the action literature and applied predictive manipulations from the broader sensory cognition literature. Experiment 1 replicated typical findings that self-produced forces are rated as less intense than similar forces presented in the absence of action. However, this effect was reversed when cutaneous stimulation was removed from the active finger. Experiments 2 and 3 manipulated the predictability of tactile action outcomes and found that expected tactile action outcomes were perceived *more*, not less, intensely than those that were unexpected. Computational modelling suggested that expectations alter the way sensory evidence is integrated – increasing the gain afforded to expected tactile signals.

These findings are consistent with upweighting perceptual accounts from outside of action domains, proposing that prior expectations are combined with sensory evidence to generate veridical perceptual interpretations of our noisy environment – thereby rendering expected events more intense. The present findings indicate that these upweighting mechanisms operate similarly in touch, and are harder to reconcile with prominent downweighting theories from action (Blakemore et al., 1998; Dogge et al., 2019; Fiehler et al., 2019; Kilteni & Ehrsson, 2017), which propose that expected action outcomes are perceptually attenuated. It is therefore essential to consider how the present findings can be resolved with the multitude of data cited in support of downweighting theories. For example, attenuating internally-generated electric fields in Mormyrid fish is thought to improve detection of prey-like stimuli (Enikolopov et al., 2018), virtual reality trained mice show suppressed auditory responses to self-produced tones generated by treadmill running (Schneider et al., 2018), and humans show reduced perceived force of tactile events during movement (Bays et al., 2005, 2006). However, studies have not demonstrated whether underlying mechanisms operate according to stimulus probabilities. There are a number of non-predictive mechanisms which could instead explain attenuation (Press et al., 2020; Press & Cook, 2015), and based upon the current findings we propose that some effects are perhaps instead generated by identity-general gating mechanisms, and others by mechanisms shaping perception according to event repetition – given that repetition is frequently confounded with expectation (Feuerriegel et al., 2020).

Importantly however, the present data should not be taken to reflect that predictive attenuation does not occur, but rather that the mechanisms generating attenuation are unlikely to operate as assumed across the last few decades. In other words, the assumed mechanisms underlying predictive tactile attenuation require pre-emptive downweighted perception of expected events – perhaps due to subtraction of the prediction from actual input (Wolpert & Flanagan, 2001) – but such a mechanism is hard to reconcile with the present findings demonstrating upweighted sensory gain of predicted action outcomes. These data more likely suggest that purported predictive mechanisms must have the capability of generating both up- and downweighting, but under different circumstances. Some of us have recently outlined how this may be achieved via opposing processes with differing roles. Our theory (Press et al., 2020) suggests that perception is initially biased towards what we expect in order to generate experiences rapidly that, on average, are more veridical. This is achieved via pre-emptively upweighting the perceptual representations of expected events. However, if events are presented which generate particularly high surprise (defined according to the Kullback Leibler Divergence; Itti & Baldi, 2009), later processes adapted to subserve learning highlight these events. Therefore, any relative downweighting of the expected is achieved via reactive processes that prioritise only the most informative of unexpected inputs. Our findings are consistent with this possibility given that tactile events were punctate (30ms) and such frequently presented unexpected events would not have generated high surprise (KLD) on average.

Regardless of the ultimate resolution of this debate, the important conclusion from the present studies is that sensorimotor prediction does not appear to exhibit a qualitatively distinct influence on tactile perception – demonstrating the need to update theoretical accounts to incorporate the vast array of data revealing both up- and downweighting across perceptual disciplines. The assumption that action predictions have a particular and distinctive attenuating influence on perception has been used to advance a number of theoretical accounts, such as those explaining sensory differences associated with healthy ageing (Wolpe et al., 2016), motor severity in Parkinson’s disease (Wolpe et al., 2018) and theories of hallucinations in clinical disorders such as schizophrenia and diseases such as Parkinson’s and Alzheimer’s (Corlett et al., 2019). However, if predictions shape perception similarly regardless of domain then these theories need revisiting.

To conclude, these findings suggest that sensorimotor prediction can increase, as well as decrease, the perceived intensity of tactile events. These findings suggest that sensorimotor prediction operates via qualitatively similar mechanisms to other prediction and regardless of the sensory domain.

## Data Availability

All data and documentation will be deposited in the Birkbeck Research Data Depository (BiRD - https://researchdata.bbk.ac.uk/) and be openly accessible by an associated DOI.

## Author Contributions

All authors contributed to developing the study concept and design. E.R.T performed data collection and analysis under the supervision of C.P. All authors approved the final version of the manuscript for submission.

## Acknowledgments

Leverhulme Trust (RPG-2016-105) and Wellcome Trust (204770/Z/16/Z) grants awarded to C.P supported this work. F.P.d.L was supported by a Vidi Grant (Nederlandse Organisatie voor Wetenschappelijk Onderzoek, 452-13-016) and ERC Starting Grant (Horizon 2020 Framework Programme, 678286).

